# In-vitro Analysis of Hemostatic Cotton Khadi Dressing with Kaolin/Chitosan-based Composition and Its Application as Prospective Wound Dressing

**DOI:** 10.1101/2022.08.12.503747

**Authors:** Aayushi A Raval, Vishwa N Patel, Utkarsh V Pancholi

## Abstract

The current investigation illustrates the formulation of Kaolin/Chitosan-based composition by using a simple mechanochemical method that can produce a stabilized hemostatic composition. Chitosan is stabilized in sodium hydroxide solution with which kaolin solution is mixed under controlled conditions. When the formulated hemostatic composition was impregnated on cotton khadi dressing it showed drastically improved hemostatic efficiency as compared to standard cotton gauze. The tightly woven cotton khadi fabric has a larger surface area as compared to standard cotton gauze and so it retains more hemostatic composition. The hemostatic properties of the composition are characterized using various in-vitro testing techniques like bleeding time analysis, clotting time analysis, absorption strength study, and several other parameters. Investigation of the freshly drawn blood samples from humans was used for experimenting in order to obtain highly accurate results. The results demonstrate that Kaolin/Chitosan composite impregnated cotton khadi dressing can noticeably enhance the hemostasis other than being biocompatible as demonstrated through an animal model and fresh blood study.

## Introduction

Globally, approximately 1.3 million people die each year due to road traffic accidents according to the World Health Organization^1^. The main cause of death in the majority of road accidents is excessive blood loss from the site of injury before any medical assistance is provided to the patient. In the majority of traumatic injuries, hemorrhage constitutes up to 40% of deaths out of which 33-56% of death occurs before the patient is hospitalized^2,3^. Hemorrhage conditions may also be encountered during outdoor activities such as trekking, camping, or adventure sports. During such traumatic conditions, hemostasis plays the most important part in protecting the body.

In daily life, the basic knowledge of first-aid suggests covering the site of bleeding using a clean piece of cloth, patch, or gauze and applying direct pressure on the site to stop the bleeding. But in traumatic conditions when the blood flow is severe, conventional first-aid treatment is not adequate to stop the blood flow and facilitate the body’s natural hemostasis. The rapid hemostasis during the injury plays a vital role in avoiding fatality due to excessive blood loss. This inspires the development of advanced and effective hemostatic dressings for prehospital wound dressing to increase the survival rate ^4^. Notably, there are various hemostatic materials and technologies developed in the market to actively facilitate the coagulation of blood at the site of injury^4–7^. Using these materials there are different types of hemostatic dressings developed for use in traumatic injury and actively coagulate the site of bleeding^8^. The main function of the hemostatic dressing is: (i) to concentrate the blood cells at the site of hemostasis and absorb the water content from the blood (ii) to rapidly activate the coagulation cascade for faster coagulation and (iii) to form a mechanical barrier and stop the bleeding ^4,7,9^. An ideal hemostatic dressing for traumatic injuries and prehospital application must be quick and easy to use, it is very much useful if it is self-applicable. It must be safe, effective, and biocompatible and it should have an easy removal procedure^10^.

Based on these parameters various hemostatic dressings are available in the market. Kaolin clay is a naturally occurring cation-based aluminosilicate that activates the intrinsic pathway of coagulation by concentrating the clotting factors at the site of bleeding and rapidly absorbing the water content from the blood^11,12^. The Quikclot combat gauze (by Z-Medica) is a widely used hemostatic dressing that is impregnated with kaolin clay^5,9^. Quikclot Combat gauze facilitates rapid coagulation of the bleeding site and is a widely used hemostatic dressing among US Army and civilians for rapid hemostasis^13^. Several hemostatic dressings with kaolin and a combination of other chemicals are available in the market^14–17^. However, the presence of only the fine clay particles sometimes poses a risk of thrombosis which may arise due to the particle detachment from gauze during use. The presence of clay reduces the mechanical strength and flexibility of the gauze and it also fails to achieve rapid hemostasis due to the high loss of active particles^18^. Chitosan occurs naturally in the shells and outer skeleton of shrimp, crabs, and other living organisms. Chitosan is a polysaccharide composed of D-glucosamine which degrades in the body and forms glucosamine and N-acetyl glucosamine^19^. Chitosan is extensively used in the pharmaceutical industry because of its antimicrobial, antioxidant, and bio-adhesive properties. Chitosan is highly biocompatible and has a major application in hemostatic dressing^20^. A previous study showed that chitosan has a positive charge which induces the aggregation of platelets and erythrocytes. It also triggers the contact system activation when it comes in contact with the material^21^. Despite all the favorable properties of chitosan for its application in hemostatic dressing, the most important concern is its solubility. Chitosan has an amino group which makes it a weak base and so it is insoluble in water. This limits the preparation methodology and combines it with other hemostatic agents.

For the testing and characterization of the hemostatic composition, various choices are available, but non-woven cotton gauze is conventionally used^13,14^. However, in this study, we have used cotton khadi dressing for its superior biocompatible properties to the standard non-woven cotton gauze. Khadi is a hand-woven or hand-spun natural fiber of cotton, silk, or woollen fibers. Among all the different combinations of khadi fabric, cotton khadi is the most versatile fabric for dressing applications. Cotton khadi is highly absorbent and thermally conductive than any other khadi fabric^22^. ^23^As the cotton fibers are natural, the cotton khadi dressing is highly biocompatible making it suitable for the invitro dressing applications.

The current study aims at formulating a stable Kaolin – Chitosan-based composition by using a mechanochemical method that can produce a stabilized hemostatic composition that can control the severe bleeding in traumatic conditions of external wounds. The cotton khadi gauze is used as the base (i.e., carrier for hemostatic composition) for coating the composition and facilitating various testing on it. The formulated composition is tested by various in-vitro testing models to characterize the efficiency of the composition. The successfully formulated composition of Kaolin/Chitosan will significantly promote blood clotting and reduce the bleeding time to a greater extent as compared to standard dressing without hemostatic composition coated on it. The cotton khadi fabric as a carrier of the hemostatic composition is a highly absorbent, biocompatible, and eco-friendly material for wound dressing.

## Materials and Methodology

### Preparation of chitosan – kaolin-based hemostatic composition

In preparation of the mixture, the two hemostatic agents (kaolin, and chitosan) are used. Kaolin is an aluminosilicate clay that is abundantly available in the crust of the earth. Kaolin is also called ‘healing clay’ because of its various healing properties^24^. Kaolin clay particles have a stacked arrangement of layers and the structure is very simple with zero net electric charges. When kaolin comes in contact with water molecules these bonds break down and allow the stacked surfaces to slip away and disperse in the liquid which makes it partially water-soluble. Kaolin is partially soluble in acidic solutions. In this study Kaolin, extra pure is acquired from Loba Chemie, and Laboratory reagents and fine chemicals are used. Kaolin particles of sizes ranging from 3-8 µm are used for the formation of hemostatic composition. The kaolin powder was filtered using gravity filtration before use. Filtered kaolin is gradually added to distilled water with continuous heating at 65C and stirring to form the milky white solution.

Chitosan occurs in abundance in nature and is an alkaline polysaccharide that facilitates rapid blood coagulation and wound healing. In this study, 75% - deacetylated chitosan (Poly (D-glucosamine)) extracted from shrimp cells is acquired from Loba Chemie (Loba Chemie, Laboratory reagents, and fine chemicals) is used. This chitosan is in the form of fibers and is insoluble in water. The dissolution of chitosan has always been a limitation for its application not being soluble in water. Chitosan is partially soluble in acidic solvents depending on the concentration of the acid the rate of dissolving of chitosan increases^25^. In this study, chitosan is dissolved in 0.1M acetic acid solution with continuous heating and stirring for 3-5 hrs. The method to coat the cotton gauze in a heated kaolin solution is conventionally used^26,27^. However, one study result shows that direct coating of kaolin slurry results in low coating efficiency and gives relatively low hemostatic efficiency^28^. To achieve higher hemostatic efficiency and lesser detachment of kaolin from the dressing the Kaolin solution is gradually added to the chitosan solution with continuous heating and stirring the solution. Chitosan solution is slightly sticky and disperses kaolin particles evenly and traps them in the solution. As the acetic acid is used in the solution to dissolve the chitosan the acidity of the solution is very high with pH 1.8. To neutralize the solution 0.1M NaOH solution is added to the hemostatic composition with continuous vigorous stirring. This process yields milky white kaolin-chitosan composition with water-like fluidity and minor sticky texture. It has been studied previously that the mixture of kaolin particles is very well dispersed in the chitosan matrix and there are strong interactions between them as disclosed in FTIR and thermogravimetric analysis^29^ The total amount of hemostatic agents used is ca. 65% of the total weight of hemostatic dressing.

### Impregnating kaolin-chitosan hemostatic dressing on cotton khadi fabric

A highly absorbent cotton khadi gauze is used as a carrier for the kaolin-chitosan hemostatic composition developed. Hand-woven khadi gauze can absorb and retain more composition as compared to non-woven standard cotton gauze. In this study, a 6-ply cotton khadi gauze procured from Khadi Gram Udyog India with a yarn ratio (warp x weft) of 12:12 is used. The khadi gauze is wetted in the hemostatic composition and the solution is allowed to be absorbed through the gauze. The impregnated gauze is heated in an oven to remove the water content from it. The final khadi dressing is sterilized in an autoclave. The kaolin and chitosan are homogenously impregnated throughout the gauze as confirmed by microscopic examination. Mixing of the kaolin and chitosan has also been accomplished as they formed an even, homogenous mixture without lumps, amalgamation, or precipitation of any component.

## 3. Results and discussion

### 3.1) In-vitro Assessment in Animals

The research protocol was revised and approved by the ‘Institutional Animal Ethics Committee (IAEC)’ appointed by ‘The Committee for the Purpose of Control and Supervision of Experiments on Animals (CPCSEA)’ India. All the albino rats received proper care and were used in strict compliance with the Guide for the Care and Use of Laboratory Animals.

#### 3.1.1) In vitro hemostatic efficiency in Rats

Ten adult albino rats were approved by the Institutional Animal Ethics Committee (IAEC) for the study and the rats were taken for experimentation from the Animal House at the Department of Pharmacology, LM College of Pharmacy, Ahmedabad, India. The adult albino rats (males and females) weighed ca. 2 kg. The rats were kept in big cages at room temperature and were allowed food and water ad libitum. Before the experiment, the rats were given mild anesthesia by inhalation of isoflurane using the bell jar technique for about 3-4 minutes. The hair of the rat at the site of experimentation was shaved using a sterile blade and razor for clear examination and application of the gauze at the site of bleeding. For the experiment rats were randomly selected and divided into groups. Among each group, one rat served as a control group and the other rats received the hemostatic khadi dressing. All the rats were maintained under anesthesia throughout the experiment and were given treatment until the wound was healed. The rat model for tail puncture was developed using a disposable 18-gauge needle. The surface area of the tail is cleansed with alcohol before inserting the needle. The first rat is used as a control group that received the standard cotton gauze. The second and third rats received Hemostatic khadi dressing. After the puncture of ca. 1 cm deep, the hemostatic khadi dressing was immediately applied and the bleeding time was examined by removing and reapplying the gauze at the interval of 10 sec. During the experiment, no surgical procedure or fluid resuscitation was allowed. The initial weight of the hemostatic khadi dressing before the experiment and after the experiment was noted to measure the amount of blood loss. The same procedure was followed as described for ca. 1 cm deep leg vein puncture and ca. 1 cm deep and for 2 cm wide dorsal skin incision.

As shown in Figure (2) the hemostatic efficiency of the standard gauze is very less as compared to the hemostatic khadi dressing. The mixture of two hemostatic components in a dressing effectively reduces the clotting time as compared to the gauze without a hemostatic dressing. In various models of puncture and incision, the khadi hemostatic dressing performed relatively faster and more effectively in blood coagulation. Kaolin rapidly promotes blood coagulation by activating the factor XII protein which initiates the coagulation cascade^30,31^. In a previous study, it has been analyzed that the kaolin/chitosan composites showed significantly reduced clotting time and enhanced hemostatic efficiency^28^. The data and results of the study are in complete agreement with what we have observed in an animal model and freshly drawn human blood samples.

**Figure 1.**
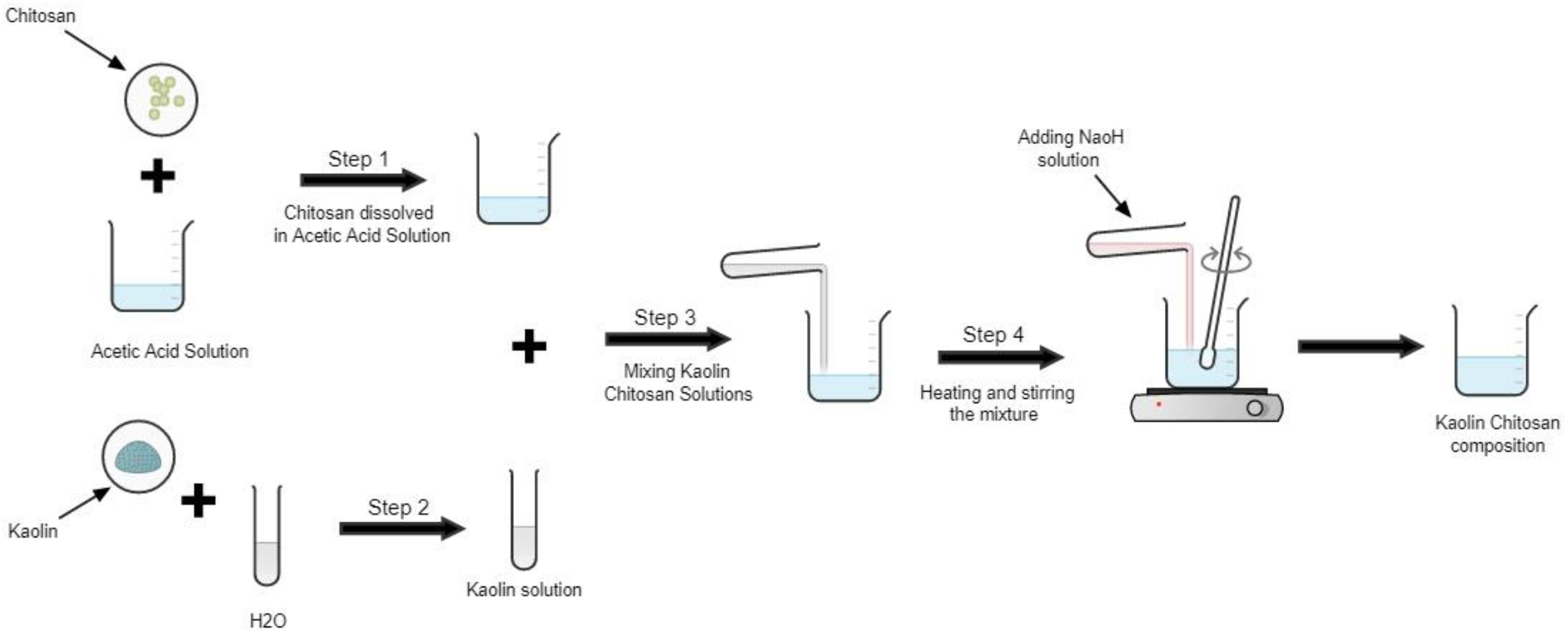
Method of preparation of kaolin/chitosan hemostatic composition

**Figure 2.**
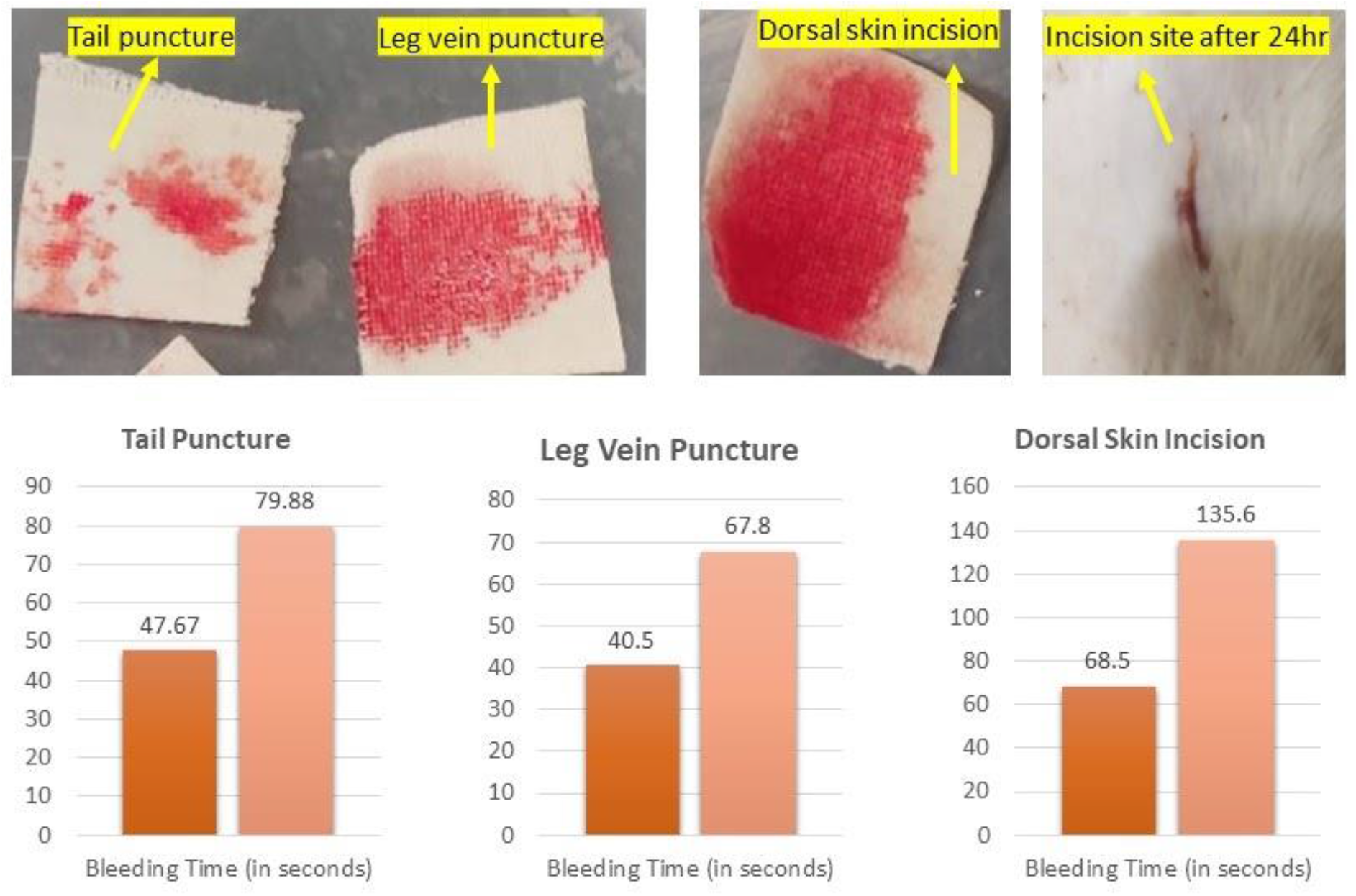
Bottom (From Left) (i) Average of the bleeding time (in seconds) calculated in two rats after puncturing their tail and reapplying hemostatic dressing every 10 seconds until the time when the bleeding stopped. (ii) Average of the bleeding time (in seconds) calculated in two rats after puncturing their leg vein and reapplying hemostatic dressing every 10 seconds until the time when the bleeding stopped. (iii) Average of the bleeding time (in seconds) calculated in two rats after puncturing their leg vein and reapplying hemostatic dressing every 10 seconds until the time when the bleeding stopped. In all the models the respective dressings were applied immediately after the injury with immediate compression.

**Figure 3.**
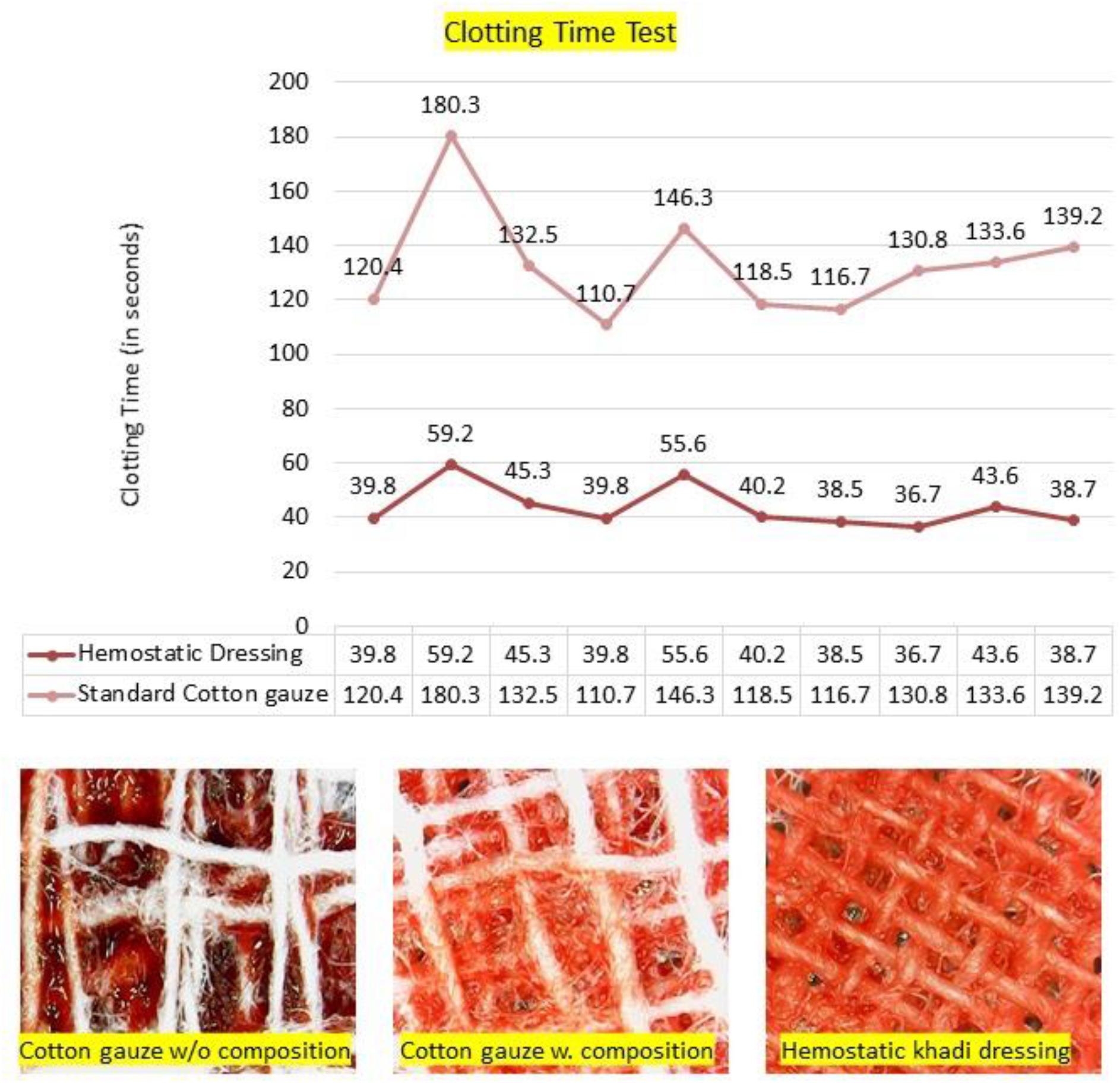
Average clotting time is calculated after the 30µl of blood comes in contact with the gauze to the time it is fully clotted. The comparative analysis shows that the standard cotton gauze requires an average of 132.9 seconds while the khadi gauze impregnated with kaolin/chitosan hemostatic agents clots the blood in 43.7 seconds. The difference between the clotting time of the standard cotton gauze and khadi hemostatic gauze is significantly large and from this, it can be noted that the khadi hemostatic agent significantly promotes blood clotting and reduces the clotting time. Figure 4. (From Left) (i) A 30µl of blood on standard cotton gauze without hemostatic dressing after 45 seconds shows no blood clotting. (ii) A 30µl of blood on standard cotton gauze with a hemostatic dressing after 45 seconds shows the onset of clotting but it is not able to significantly absorb the blood. (iii) A 30µl of blood on khadi hemostatic dressing after 45 seconds shows complete blood clotting and it can significantly absorb the blood and promote hemostasis.

**Figure 3.**
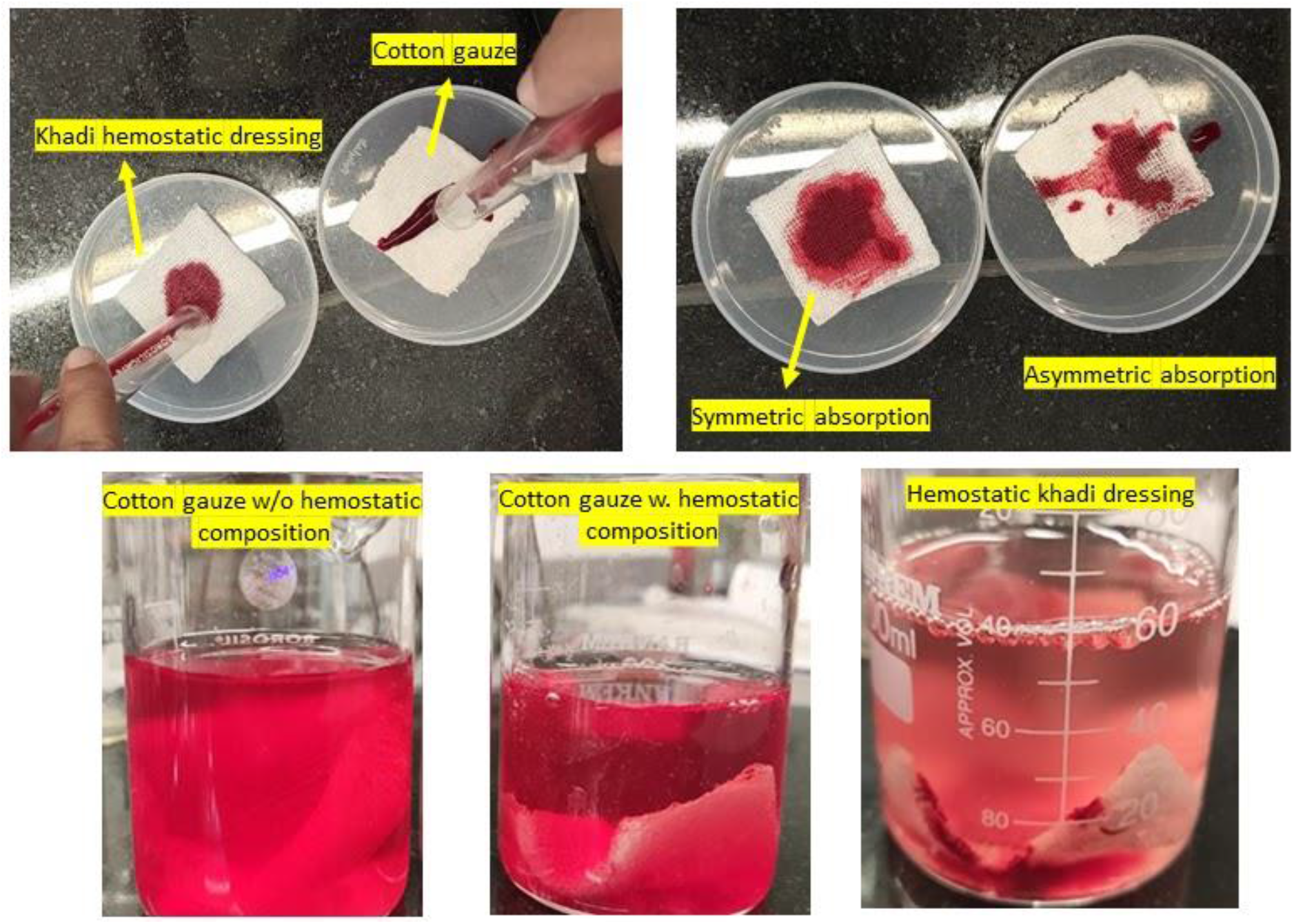
Top (From Left) (i) Blood sample of 1mL of blood is poured on khadi hemostatic gauze (first from left) and standard cotton gauze (second from left) at time t=0 second simultaneously. (ii) The absorption rates of both the dressings are observed and after 30 seconds from the initial time (t=0 seconds), an image is captured for comparative clotting and absorption analysis. The khadi hemostatic dressing performs significantly better as compared to the standard cotton gauze in absorption and clotting the blood sample. Bottom (From left) (i) The standard cotton gauze with no hemostatic agents impregnated on it was dropped into the beaker and after 60 seconds most of the blood on the gauze was dissolved in water as can be seen from the bright red color of the solution. (b) The standard cotton gauze with hemostatic agents impregnated on it was dropped in the beaker and after 60 seconds some of the blood on the gauze was dissolved in water as can be seen from the light bright red color of the solution. As compared to the first gauze the water is slightly clear which suggests that the presence of hemostatic agents has improved the clot formation, however, the strength of the clot is weaker. (iii) The khadi hemostatic dressing was dropped in the beaker and after 60 seconds very little amount of the blood on the gauze was dissolved in water as can be seen from the light pink color of the solution. As compared to the first and the second gauze the water is very clear which suggests that the presence of hemostatic agents has highly facilitated the clot formation and the strength of the clot is very stronger because of the woven structure of the khadi.

#### 3.1.2) In vitro Biocompatibility Examination

The animal model was developed by dorsal skin incision and immediately applying hemostatic khadi dressing with light manual pressure. All the adult albino rats were maintained under general mild anesthesia during the application of khadi hemostatic dressing. After the clotting, the hemostatic khadi dressing was applied to the dorsal skin and was kept for 24 hours. The dorsal skin was checked every 5 hours to check the changes if any. The animal did not show any allergies, discoloration at the site, pigmentation of the skin, or any other visual changes at the site of the injury. The animal did not show any signs of fever or allergy. This study clearly shows that the application of khadi hemostatic dressing is safe and biocompatible for in vitro applications. The dressing was easily removed with saline after 24 hours and the animal was allowed to heal naturally by applying soframycin on the incision. The animal recovered the incision completely in 5-7 days. During recovery, the animal was kept in close observation in a separate cage at room temperature and it was allowed food and water. During the recovery duration, it did not show any allergies, discoloration of the skin, pigmentation of the skin, or any other visual change at the site. The animal did not show any fever or weight loss during recovery. The hemostatic khadi dressing demonstrated the biocompatibility of the hemostatic khadi dressing after 24 hrs of continuous application on the dorsal skin incision. After 24 hrs the dressing was easily removable with saline. In conclusion, in the rat model with different types of injury model, hemostasis has been achieved on applying the hemostatic khadi dressing whereas significantly higher blood loss and longer clotting time were observed after utilizing the standard cotton gauze.

### 3.2) In-vitro hemostatic efficiency in humans

The research protocol was reviewed and approved by the “Diagnostics Research Laboratory” by the faculty of the Biomedical Department in L.D. College of Engineering, Ahmedabad, India. All the volunteers who volunteered for donating very small quantities of blood have explained the procedure and the experiment before withdrawing the blood. The blood was collected after the consent of the donor by a trained practitioner and under the supervision of in charge of the experiment.

#### 3.2.1) In vitro hemostatic efficiency in freshly drawn human blood

Ten people (both male and female) of the age group between 18-60 years volunteered for donating ca. 2ml of blood for testing the efficiency of khadi hemostatic dressing. Before the experiment, the volunteers were informed about the process of blood extraction and its use in the experiment. After their consent, the blood was collected by trained laboratory personnel using a 24-gauge x 1 TW disposable syringe. The blood was collected by vein puncture from the cephalic vein, median cubital vein, or basilic vein of the arm depending on the visibility. Each volunteer donated ca. 2ml blood one time which was utilized for the testing. The experiment setup was prepared prior to the blood collection to immediately use the freshly drawn blood and minimize the observation error. The blood collected is stored in an Eppendorf tube with heparin in it to prevent blood coagulation.

The total volume of blood collected each time (i.e., ca. 2ml) was divided into two groups. Half-blood volume (i.e., 1ml) was used as a control group for testing on the standard cotton gauze. The other half-blood volume was used for testing on the khadi hemostatic dressing. The experiment setup was done prior to blood collection and after blood collection, multiple observations were taken the experiment. Using a micropipette, 30µL of freshly drawn blood was used for clotting time analysis on standard cotton gauze as the control center and khadi hemostatic gauze. The time was noted and the observation of blood clotting was done manually using a digital microscope. The dressing without the hemostatic agent performed poorly as compared to the dressing with the hemostatic agents impregnated. The hemostatic khadi dressing reduced the clotting time to about half that of the one with the standard cotton gauze. The difference in blood clotting time between standard cotton gauze and khadi hemostatic gauze is statistically significant.

#### 3.2.2) Comparative analysis of hemostatic efficiency of dressing

To examine the comparative analysis the blood clotting experiment as mentioned in (3.2.1) it was performed on standard cotton gauze, hemostatic composition impregnated on standard cotton gauze and the hemostatic khadi dressing developed. The same procedures that have been described previously for blood clotting efficiency have been used for this experiment as well. Before beginning the experiment, blood samples were collected and the same process was performed as described in the previous experiment (3.2.1) to compare the blood clotting. The standard cotton cloth dressing without any hemostatic composition impregnated on it performed poorly as compared to cotton gauze with hemostatic agents impregnated on it and the khadi hemostatic dressing.

As the standard cotton gauze dressing is non-woven with its cotton strands loosely packed in a mesh arrangement it is not able to absorb and stop the blood like woven khadi gauze. The blood flows down through the spaces between the cotton strands and it proves ineffective in promoting blood clotting as compared to the gauze with hemostatic dressings impregnated on them. When the standard cotton gauze is coated with kaolin/chitosan composition it performs better than the cotton gauze without hemostatic agents impregnated but as the cotton is non-woven there is the blood is still able to flow down through the spaces between the cotton strands. However, the khadi cotton fabric is hand-woven and tightly packed which retains more composition as compared to standard cotton gauze and actively promotes hemostasis. As soon as the blood touches the surface of the fabric it gets absorbed and the high fiber density and closed-packed structure stop the blood loss. The comparative analysis examinations demonstrate that the khadi hemostatic dressing performed statistically better as compared to the standard cotton gauze without hemostatic composition and the standard cotton gauze with hemostatic composition impregnated. The khadi hemostatic dressing resulted in a higher blood clotting rate which can be explained by the time to hemostasis and lower amount of blood loss and lesser bleeding time.

### 3.3) Characterization of hemostatic efficiency

The research protocol was reviewed and approved by the “Diagnostics Research Laboratory” by the faculty of the Biomedical Department in L.D. College of Engineering, Ahmedabad, India and by the “Pharmacology Laboratory” by the Department of Pharmacology in LM College of Pharmacy, Ahmedabad, India. All the volunteers who volunteered for donating very small quantities of blood have explained the procedure and the experiment before withdrawing the blood. The blood was collected after consent by a trained practitioner and under the supervision of in charge of the experiment.

#### 3.3.1) Characterization of blood absorption

The same procedure is followed as discussed in detail in the prior experiment (3.2.1) to collect fresh blood from the volunteer. The freshly drawn blood is used to check the absorption efficiency of the khadi hemostatic dressing. The standard cotton gauze is used as a control sample for the experiment. In order to test the absorption rate of the standard cotton gauze and khadi hemostatic dressing, 1ml of freshly drawn blood is poured simultaneously on standard cotton gauze and khadi hemostatic dressing. The time is calculated as soon as the blood is decanted on both the dressings simultaneously.

Upon visual analysis, the significant difference in absorption rate is very clear. The khadi hemostatic dressing absorbs the blood sample faster compared to standard cotton gauze. The khadi fabric strands are very closely packed as it is handwoven and so it offers a larger surface that comes in contact with the blood. The larger surface area of the fabric retains a larger amount of kaolin/chitosan composition and so it also promotes faster blood clotting. The blood sample is evenly absorbed and the clotting is faster as compared to the standard hemostatic gauze. The comparative analysis shows that the standard cotton gauze performed poorly and the blood sample takes a very long time to get absorbed in it. As the gauze is non-woven the blood sample poured on the gauze flows down through the spaces between the cotton strands and it does not facilitate in stopping blood loss or helping in blood clotting. It significantly increases the bleeding time as a very lesser surface area comes in contact with the bleeding site.

#### 3.3.2) Characterization of clot strength

To study the clot strength and compare it with the standard cotton gauze, cotton gauze with hemostatic dressing the following experiment has been conducted. Freshly drawn blood is used for the experiment and the same procedure has been used as discussed previously. The blood sample of ca. 1ml is poured on the standard cotton gauze, cotton gauze with hemostatic dressing impregnated on it, and the khadi hemostatic dressing simultaneously. The blood sample is allowed to absorb and clot without any disturbance for 45 seconds.

After 45 seconds all the gauzes were dropped in the beaker with 60ml clean water in it and were allowed to dissolve in water. After 60 seconds of dropping in the beaker, the results were observed. The standard cotton gauze with no hemostatic composition impregnated on it performed poorly, as soon as it came in contact with the water the blood was mixed with water. As it has no hemostatic agents there is no visible clotting on its surface and the clot strength is non-existent. The cotton gauze with hemostatic agents impregnated on it performed a little better as compared to the standard cotton gauze in the experiment. As the hemostatic agents are present on the surface there was some amount of clotting on the gauze, however, the non-woven structure with larger distances between two strands of fabric do not strengthen the clot formed. As soon as it comes in contact with water the blood and the clot form slowly and significantly start dissolving in water. The khadi hemostatic dressing performed significantly better as compared to both the gauzes. The khadi cotton is tightly woven and naturally has a higher absorption rate which absorbs and retains the larger amount of blood on it. The closely packed strands strengthen the blood clot and prevent it from dissolving in water. As compared to standard cotton gauze and cotton gauze with hemostatic agents impregnated, the khadi hemostatic gauze has significantly stronger clots and least dissolution in water which explains faster clotting and very less bleeding time.

**Table 1.** Comparative analysis of the visual differences among the standard cotton gauze, standard cotton gauze with hemostatic agents impregnated in it, and khadi hemostatic dressing respectively.

| Parameters | Standard Cotton gauze with no hemostatic agents | Standard Cotton gauze with hemostatic agents | Khadi hemostatic dressing |
| --- | --- | --- | --- |
| Clarity of water | Unclear | Unclear | Partially clear |
| Colour intensity of dissolved blood | Very Bright red | Bright red | Very light pink |
| Dissolution of impregnated hemostatic composition | NA | Some amount of solution was dissolved in water | Very few traces of dissolved solution were found in the water |

## Conclusion

Our aim has been to develop the cotton khadi fabric hemostatic dressing with the enhanced hemostatic property. We reported an approach to provide hemostatic ability to the khadi fabric. The closely packed woven structure of cotton khadi fabric serves as an increased functional surface area for the application of the hemostatic composition. The results confirmed that the combination of kaolin/chitosan improved the hemostatic efficiency and promoted rapid blood clotting. Moreover, the hemostatic khadi dressing exhibited biocompatibility for in-vitro use in animal models. On the other hand, the cotton khadi fabric and the kaolin/chitosan together make a highly viable and cost-effective wound dressing. The significantly improved hemostatic efficiency and biocompatibility profiles illustrate that the hemostatic khadi fabric has enormous potential for its application as an in-vitro wound dressing.

## Acknowledgment

We gratefully acknowledge the financial support from the Student Start-up Cell -SSIP sanction letter No. LDCE/SSIP/PoC_sanction/Phase2/5082-1 (Grant No: LDCE/SSIP/BME/Phase2/004), Government of Gujarat, India. We heartily thank Dr. Mrunal Chaudhry, Dr. Vishwas Shah, Utkarsh Upadhyay, Rudradutt Thaker, and Kaxan Raval for volunteering in fresh blood sample donation for the study. We sincerely thank Dr. Mahesh Chhabria, Dr. Gaurang Shah, and LM College of Pharmacy, Ahmedabad, India for very kind collaboration for the research work. We thank Dr. Vishwas Shah and Shagun Sir for their constant help and support during the animal testing.

